# Monkey Pox Virus (MPXV): Phylogenomics, Host-Pathogen Interactome, and Mutational Cascade

**DOI:** 10.1101/2022.07.25.501367

**Authors:** Roshan kumar, Shekhar Nagar, Shazia Haider, Utkarsh Sood, Kalaiarasan Ponnusamy, Gauri Garg Dhingra, Shailly Anand, Ankita Dua, Mona Singh, Manisha Sengar, Indrakant Kumar Singh, Rup Lal

## Abstract

While the world is still managing to recover from Covid-19 pandemic, Monkeypox awaits to bring in another global outbreak as a challenge to the entire mankind. However, Covid-19 pandemic have taught us lessons to move fast in viral genomic research to implement prevention and treatment strategies. One of the important aspects in Monkeypox virus should be immediately taken up is to gather insights of its evolutionary lineage based on the genomic studies. We have thus analysed the genome sequences of reported isolates of Monkeypox in the present study through phylogenomics. Host-pathogen interactions, mutation prevalence and evolutionary dynamics of this virus were investigated for all the documented isolates. Phylogenetic exploration revealed the clustering of strain Israel 2018 (MN 648051.1) from Clade I with the four isolates reported from the recent outbreak. An in-depth scrutiny of the host-pathogen interactome identified protein E3, serine protease inhibitor-2 (SPI-2), protein K7, and cytokine response-modifying protein B (CrmB) as the major regulatory hubs. Among these, the CrmB protein (dN/dS ≈ 1.61) was detected to be operating through positive selection. It possibly attests a selective advantage with the monkeypox virus in protecting the infected cells from antiviral responses elicited by the host. Studies also revealed that CrmB protein exhibited several mutations, the majority of which were destabilizing (ΔΔG >0). While this study identified a large number of mutations within the newly outbreak clade, it also reflected that we need to move fast with the genomic analysis of the newly detected strains from around the world to develop better prevention and treatment methods

## Introduction

The discovery of the first monkeypox virus (MPXV) dates back to 1959 in cynomolgus monkeys during a transit from Singapore to Denmark [1]. However, in human history the first incidence of monkeypox was reported in a 9-month unvaccinated infant in 1970 in Basankusu Territory, Democratic Republic of Congo, Central Africa [1]. Monkeypox virus (MPXV) is a double stranded DNA virus that belongs to the family Poxviridae, subfamily Chordopoxvirinae, and the genus ‘Orthopoxvirus’, and is a close relative of Vaccinia and Cowpox viruses [1]. The genus Orthopoxvirus includes viruses such as the Vaccinia virus, Cowpox virus, Camelpox virus, Rabbitpox virus, Horsepox virus, Ectromelia virus, Variola virus, Buffalopox virus, Akhmeta virus, etc, to name a few. Human monkeypox is a self-limiting, viral zoonotic disease found mostly in people residing in areas adjacent to the tropical rainforests, and is mainly transmitted through blood, body lesions, and fluids of infected animals, shedding of viral particles through faeces and sharing contaminated items [2]. According to World Health Organization (WHO) guidelines, the symptoms usually manifest within 6 to 13 days post infection but in some cases, the incubation may even last between 5 to 21 days. The infection can present severe symptoms especially in young children, pregnant women and immunosuppressed individuals where the fatality rate can even rise to 3-6% (https://www.who.int/news-room/fact-sheets/detail/monkeypox). Clinical manifestations are similar to smallpox as the infection is presented by fever, headache, muscle ache, backache, rashes all over the body and lymphadenopathy (swelling of lymph nodes) [3]. The enlargement of lymph nodes is exclusively observed in patients with MPXV infection and helps differentiate it from smallpox symptoms. Additionally, MPXV has been designated less severe than smallpox with a lower mortality rate.

Geographically, the disease has been endemic to central and western Africa, however, in the past few years, reports of human-human and nosocomial transmission have emerged that can make MPXV another potential global threat reeling under post COVID complications. In the United Kingdom, MPX has been named as High Consequence Infectious Disease (HCID) and special facilities are maintained to treat such patients [3, 4]. On July 23, 2022, the World Health Organisation declared Monkeypox Public Health Emergency of International Concern (PHEIC). The incidence was presumed to be associated with the import of Gambian giant rats, squirrels and dormice that transmitted the virus to prairie dogs domesticated as pets [5]. Recently, this pathogen has resurfaced and as of July 22, 2022, according to the reports of the European Centre for Disease Prevention and Control, more than 18000 cases of monkeypox have been confirmed in different parts of the world including Spain, USA, Germany, United Kingdom, France and India. The sudden emergence of MPX and its widespread prevalence in more than 70 locations indicate that the virus must have been prevalent and been circulating at levels that have gone undetected by surveillance systems. Realizing this. As the spread has been attributed to travel and numbers are slowly increasing globally, the same cardinal mistakes must not be repeated and screenings of international travellers must be more rigorous this time to prevent multi-country outbreaks.

In order to tackle the outbreak of Monkeypox we need to move fast on several fronts and one of the aspects that really helped scientists and policy makers on case of SARS-Covid - 2 pandemic was very rapid genomic analysis of strains from different geographical locations [6, 7]. Indeed, pathogen genomic studies have been very helpful in characterizing different circulating strains, identifying evolutionary links, and predicting transmission patterns. In the present study we thus attempted to analyse the phylogenetic position of the newly emerged sequenced Monkeypox virus variants within the genus Orthopoxvirus. We particularly focused on the genomic variation present at protein levels within monkeypox strains followed by the generation of an interactome between human and monkeypox virus. In addition, the amino acid level mutations in major regulatory hub proteins were analysed to determine their effect on virus structural integrity. Although results reflect that Monkeypox virus is harbour a lot of mutations, more in-depth analysis is needed as more and more genome sequences of this strain become available in near future to supplement the efforts to devise prevention and treatment strategies

## Materials and Methods

### Selection of Genomes

All the monkeypox virus genomes were retrieved from the NCBI database. Following retrieval of the sequences, quality assessment was performed using QUAST 5.0.2 [8] and out of the 126 genomes available, 71 high quality genomes were selected for downstream analysis including the four isolates of the year 2022 (Table 1). In order to establish the phylogenetic position, using the whole genome Single Nucleotide Polymorphism (SNP) method, all available genomes of genus Orthopoxvirus other than Monkeypox viruses were downloaded (N=100). These comprised the sequences from Cowpox viruses (N=41), Vaccinia viruses (N=30), Camelpox viruses (N=9), Akhmeta viruses (N=6), Variola viruses (N=5) and Ectromelia viruses (N=3). After the quality check, out of these 100 genomes, 85 were used for SNP-based phylogeny. Finally, the phylogenetic analysis was constructed using 156 viruses, 71 genomes of MPXV and the remaining 85 genomes of viruses belonging to the genus Orthopoxvirus.

**Table 1:** The general genomic attributes of monkey pox viruses.

| **S. No.** | **Isolate** | **Accession Number** | **Genome Size (bp)** | **CDS** | **GC%** | **Host** | **Country** | **Collection Date** | **Date of Sequence Submission** | **Publication** |
| --- | --- | --- | --- | --- | --- | --- | --- | --- | --- | --- |
| 1. | Monkeypox virus strain 015c | MN702448 | 189838 | 177 | 32.9 | Homo sapiens | Central African Republic | 07-Mar-18 | 29-Aug-20 | Unpublished |
| 2. | Monkeypox virus strain 18 | MN702447 | 189841 | 179 | 33 | Homo sapiens | Central African Republic | 20-Mar-18 | 29-Aug-20 | Unpublished |
| 3. | Monkeypox virus strain 38c | MN702446 | 190357 | 195 | 33 | Homo sapiens | Central African Republic | 14-Apr-18 | 29-Aug-20 | Unpublished |
| 4. | Monkeypox virus strain A1 | MN702453 | 189632 | 182 | 33 | Homo sapiens | Central African Republic | 11-Aug-01 | 29-Aug-20 | Unpublished |
| 5. | Monkeypox virus strain A2 | MN702452 | 190244 | 183 | 33 | Homo sapiens | Central African Republic | 10-Aug-10 | 29-Aug-20 | Unpublished |
| 6. | Monkeypox virus strain A4 | MN702445 | 190519 | 170 | 33 | Homo sapiens | Central African Republic | 06-Feb-17 | 29-Aug-20 | Unpublished |
| 7. | Monkeypox virus strain A5 | MN702444 | 190167 | 170 | 33 | Homo sapiens | Central African Republic | 06-Feb-17 | 29-Aug-20 | Unpublished |
| 8. | Monkeypox virus strain A6 | MN702451 | 190413 | 191 | 33 | Homo sapiens | Central African Republic | 14-Apr-17 | 29-Aug-20 | Unpublished |
| 9. | Monkeypox virus strain B1 | MN702450 | 190264 | 191 | 33 | Homo sapiens | Central African Republic | 01-Jan-16 | 29-Aug-20 | Unpublished |
| 10. | Monkeypox virus strain B2 | MN702449 | 190270 | 190 | 33 | Homo sapiens | Central African Republic | 02-01-2016 | 29-Aug-20 | Unpublished |
| 11. | Monkeypox virus strain Boende_DRC_2008 | KP849469 | 197422 | 188 | 33.1 | NA | Democratic Republic of the Congo | 2008 | 13-May-15 | Viruses 7 (4), 2168-2184 (2015) |
| 12. | Monkeypox virus strain Cameroon-1990 | KJ642618 | 194363 | 177 | 33.1 | NA | Republic of Cameroon | 1990 | 11-May-15 | Viruses 7 (4), 2168-2184 (2015) |
| 13. | Monkeypox virus strain Congo_2003_358 | DQ011154 | 197195 | 200 | 33.1 | Homo sapiens | NA | NA | 28-Sep-05 | J. Gen. Virol. 86 (PT 10), 2661-2672 (2005) |
| 14. | Monkeypox virus strain Congo_2003_358 | KJ642613 | 197195 | 178 | 33.1 | Homo sapiens | Zaire, Democratic Republic of the Congo | 1970 | 28-Sep-05 | Viruses 7 (4), 2168-2184 (2015) |
| 15. | Monkeypox virus strain COP-58 | AY753185 | 199469 | 177 | 33.1 | NA | NA | NA | 07-Sep-05 | Virology 340 (1), 46-63 (2005) |
| 16. | Monkeypox virus strain Cote d'Ivoire_1971 | KP849470 | 200397 | 185 | 33.1 | NA | NA | 1971 | 13-May-15 | Viruses 7 (4), 2168-2184 (2015) |
| 17. | Monkeypox virus strain DRC 06-0950 | JX878407 | 196440 | 190 | 33.1 | Homo sapiens | Democratic Republic of the Congo | 09-10-2006 | 14-Jan-14 | Emerging Infect. Dis. 20 (2), 232-239 (2014) |
| 18. | Monkeypox virus strain DRC 06-0970 | JX878408 | 196740 | 191 | 33.1 | Homo sapiens | Democratic Republic of the Congo | 31-10-2006 | 14-Jan-14 | Emerging Infect. Dis. 20 (2), 232-239 (2014) |
| 19. | Monkeypox virus strain DRC 06-0999 | JX878409 | 198597 | 191 | 33.1 | Homo sapiens | Democratic Republic of the Congo | 09-11-2006 | 14-Jan-14 | Emerging Infect. Dis. 20 (2), 232-239 (2014) |
| 20. | Monkeypox virus strain DRC 06-1070 | JX878410 | 198886 | 191 | 33.1 | Homo sapiens | Democratic Republic of the Congo | 24-11-2006 | 14-Jan-14 | Emerging Infect. Dis. 20 (2), 232-239 (2014) |
| 21. | Monkeypox virus strain DRC 06-1075 | JX878411 | 198877 | 191 | 33.1 | Homo sapiens | Democratic Republic of the Congo | 30-11-2006 | 14-Jan-14 | Emerging Infect. Dis. 20 (2), 232-239 (2014) |
| 22. | Monkeypox virus strain DRC 06-1076 | JX878412 | 198737 | 191 | 33.1 | Homo sapiens | Democratic Republic of the Congo | 30-11-2006 | 14-Jan-14 | Emerging Infect. Dis. 20 (2), 232-239 (2014) |
| 23. | Monkeypox virus strain DRC 07-0045 | JX878413 | 197627 | 191 | 33.1 | Homo sapiens | Democratic Republic of the Congo | 14-12-2006 | 14-Jan-14 | Emerging Infect. Dis. 20 (2), 232-239 (2014) |
| 24. | Monkeypox virus strain DRC 07-0046 | JX878414 | 197910 | 191 | 33.1 | Homo sapiens | Democratic Republic of the Congo | 14-12-2006 | 14-Jan-14 | Emerging Infect. Dis. 20 (2), 232-239 (2014) |
| 25. | Monkeypox virus strain DRC 07-0092 | JX878415 | 197488 | 191 | 33.1 | Homo sapiens | Democratic Republic of the Congo | 26-12-2006 | 14-Jan-14 | Emerging Infect. Dis. 20 (2), 232-239 (2014) |
| 26. | Monkeypox virus strain DRC 07-0093 | JX878416 | 197632 | 191 | 33.1 | Homo sapiens | Democratic Republic of the Congo | 26-12-2006 | 14-Jan-14 | Emerging Infect. Dis. 20 (2), 232-239 (2014) |
| 27. | Monkeypox virus strain DRC 07-0104 | JX878417 | 197959 | 191 | 33.1 | Homo sapiens | Democratic Republic of the Congo | 14-12-2006 | 14-Jan-14 | Emerging Infect. Dis. 20 (2), 232-239 (2014) |
| 28. | Monkeypox virus strain DRC 07-0120 | JX878418 | 196740 | 191 | 33.1 | Homo sapiens | Democratic Republic of the Congo | 02-01-2007 | 14-Jan-14 | Emerging Infect. Dis. 20 (2), 232-239 (2014) |
| 29. | Monkeypox virus strain DRC 07-0275 | JX878419 | 196732 | 191 | 33.1 | Homo sapiens | Democratic Republic of the Congo | 10-02-2007 | 14-Jan-14 | Emerging Infect. Dis. 20 (2), 232-239 (2014) |
| 30. | Monkeypox virus strain DRC 07-0283 | JX878420 | 196730 | 191 | 33.1 | Homo sapiens | Democratic Republic of the Congo | 26-02-2007 | 14-Jan-14 | Emerging Infect. Dis. 20 (2), 232-239 (2014) |
| 31. | Monkeypox virus strain DRC 07-0286 | JX878421 | 197767 | 191 | 33.1 | Homo sapiens | Democratic Republic of the Congo | 22-03-2007 | 14-Jan-14 | Emerging Infect. Dis. 20 (2), 232-239 (2014) |
| 32. | Monkeypox virus strain DRC 07-0287 | JX878422 | 197346 | 191 | 33.1 | Homo sapiens | Democratic Republic of the Congo | 20-03-2007 | 14-Jan-14 | Emerging Infect. Dis. 20 (2), 232-239 (2014) |
| 33. | Monkeypox virus strain DRC 07-0337 | JX878423 | 196581 | 190 | 33.1 | Homo sapiens | Democratic Republic of the Congo | 25-03-2007 | 14-Jan-14 | Emerging Infect. Dis. 20 (2), 232-239 (2014) |
| 34. | Monkeypox virus strain DRC 07-0338 | JX878424 | 196376 | 190 | 33.1 | Homo sapiens | Democratic Republic of the Congo | 25-03-2007 | 14-Jan-14 | Emerging Infect. Dis. 20 (2), 232-239 (2014) |
| 35. | Monkeypox virus strain DRC 07-0354 | JX878425 | 197147 | 190 | 33.1 | Homo sapiens | Democratic Republic of the Congo | 20-04-2007 | 14-Jan-14 | Emerging Infect. Dis. 20 (2), 232-239 (2014) |
| 36. | Monkeypox virus strain DRC 07-0450 | JX878426 | 196747 | 191 | 33.1 | Homo sapiens | Democratic Republic of the Congo | 27-05-2007 | 14-Jan-14 | Emerging Infect. Dis. 20 (2), 232-239 (2014) |
| 37. | Monkeypox virus strain DRC 07-0480 | JX878427 | 197347 | 191 | 33.1 | Homo sapiens | Democratic Republic of the Congo | 25-05-2007 | 14-Jan-14 | Emerging Infect. Dis. 20 (2), 232-239 (2014) |
| 38. | Monkeypox virus strain DRC 07-0514 | JX878428 | 197488 | 191 | 33.1 | Homo sapiens | Democratic Republic of the Congo | 30-06-2007 | 14-Jan-14 | Emerging Infect. Dis. 20 (2), 232-239 (2014) |
| 39. | Monkeypox virus strain DRC 07-0662 | JX878429 | 196866 | 190 | 33.1 | Homo sapiens | Democratic Republic of the Congo | 04-09-2007 | 14-Jan-14 | Emerging Infect. Dis. 20 (2), 232-239 (2014) |
| 40. | Monkeypox virus strain DRC Yandongi 1985 | KC257460 | 196487 | 198 | 33.1 | Homo sapiens | Democratic Republic of the Congo: Yandongi | 1985 | 20-Feb-13 | Emerging Infect. Dis. 19 (2), 237-245 (2013) |
| 41. | Monkeypox virus strain Gabon-1988 | KJ642619 | 196546 | 177 | 33.1 | NA | Gabon | 1988 | 11-May-15 | Viruses 7 (4), 2168-2184 (2015) |
| 42. | Monkeypox virus strain Ikubi | KJ642612 | 194744 | 177 | 33.1 | NA | Zaire, Democratic Republic of the Congo | 1986 | 11-May-15 | Viruses 7 (4), 2168-2184 (2015) |
| 43. | Monkeypox virus strain Israel_2018 | MN648051 | 197417 | 213 | 33 | Homo sapiens | Israel | 04-Oct-18 | 09-Jan-20 | NA |
| 44. | Monkeypox virus strain Ivory Coast 2012 | KJ136820 | 200035 | 210 | 33.1 | Wild Monkey | Cote d'Ivoire: Tai National Park | Mar-12 | 29-May-14 | Emerging Infect. Dis. 20 (6), 1009-1011 (2014) |
| 45. | Monkeypox virus strain Liberia_1970_184 | DQ011156 | 200256 | 198 | 33.1 | Homo sapiens | NA | NA | 28-Sep-05 | J. Gen. Virol. 86 (PT 10), 2661-2672 (2005) |
| 46. | Monkeypox virus strain MPXV-M2940_FCT | MT903337 | 197547 | 181 | 33 | Homo sapiens | Nigeria: FCT | NA | 15-Sep-20 | J. Infect. Dis. (2020) In press |
| 47. | Monkeypox virus strain MPXV-M2957_Lagos | MT903338 | 197559 | 181 | 33 | Homo sapiens | Nigeria: Lagos State | NA | 15-Sep-20 | J. Infect. Dis. (2020) In press |
| 48. | Monkeypox virus strain MPXV-M3021_Delta | MT903339 | 197556 | 181 | 33 | Homo sapiens | Nigeria: Delta State | NA | 15-Sep-20 | J. Infect. Dis. (2020) In press |
| 49. | Monkeypox virus strain MPXV-M5312_HM12_Rivers | MT903340 | 197209 | 181 | 33 | Homo sapiens | Nigeria: Rivers State | NA | 15-Sep-20 | J. Infect. Dis. (2020) In press |
| 50. | Monkeypox virus strain MPXV-M5320_M15_Bayelsa | MT903341 | 185309 | 182 | 33 | Homo sapiens | Nigeria: Bayelsa State | NA | 15-Sep-20 | J. Infect. Dis. (2020) In press |
| 51. | Monkeypox virus strain MPXV-Singapore | MT903342 | 197309 | 181 | 33 | Homo sapiens | Singapore | NA | 15-Sep-20 | J. Infect. Dis. (2020) In press |
| 52. | Monkeypox virus strain MPXV-USA2003_099_Dormouse | MT903347 | 198780 | 181 | 33.1 | Dormouse | USA | NA | 15-Sep-20 | J. Infect. Dis. (2020) In press |
| 53. | Monkeypox virus strain MPXV-USA2003_099_Gambian_Rat | MT903346 | 198778 | 181 | 33.1 | Cricetomys gambianus | USA | NA | 15-Sep-20 | J. Infect. Dis. (2020) In press |
| 54. | Monkeypox virus strain MPXV-USA2003_099_Rope_Squirrel | MT903348 | 198780 | 181 | 33.1 | Rope Squirrel | USA | NA | 15-Sep-20 | J. Infect. Dis. (2020) In press |
| 55. | Monkeypox virus strain MPXV-WRAIR7-61 | AY603973 | 199195 | 177 | 33.1 | NA | West Africa | NA | 02-Sep-05 | Virology 340 (1), 46-63 (2005) |
| 56. | Monkeypox virus strain Nigeria-SE-1971 | KJ642617 | 197551 | 176 | 33 | NA | Nigeria | 1971 | 11-May-15 | Viruses 7 (4), 2168-2184 (2015) |
| 57. | Monkeypox virus strain PCH | KJ642616 | 198741 | 178 | 33.1 | NA | France: Paris | 1968 | 11-May-15 | Viruses 7 (4), 2168-2184 (2015) |
| 58. | Monkeypox virus strain Sierra Leone | AY741551 | 198756 | 177 | 33.1 | NA | Sierra Leone |  | 07-Sep-05 | Virology 340 (1), 46-63 (2005) |
| 59. | Monkeypox virus strain Sudan 2005_01 | KC257459 | 206372 | 207 | 32.9 | Homo sapiens | Sudan: Nuria | 2005 | 20-Feb-13 | Emerging Infect. Dis. 19 (2), 237-245 (2013) |
| 60. | Monkeypox virus strain USA_2003_039 | DQ011157 | 198780 | 198 | 33.1 | Homo sapiens | USA | NA | 28-Sep-05 | J. Gen. Virol. 86 (PT 10), 2661-2672 (2005) |
| 61. | Monkeypox virus strain USA_2003_044 | DQ011153 | 198780 | 198 | 33.1 | Prairie Dog | USA | NA | 28-Sep-05 | J. Gen. Virol. 86 (PT 10), 2661-2672 (2005) |
| 62. | Monkeypox virus strain UTC | KJ642614 | 190083 | 172 | 33 | NA | Netherlands: Rotterdam | 1965 | 11-May-15 | Viruses 7 (4), 2168-2184 (2015) |
| 63. | Monkeypox virus strain W-Nigeria | KJ642615 | 197792 | 176 | 33 | NA | Nigeria | 1978 | 11-May-15 | Viruses 7 (4), 2168-2184 (2015) |
| 64. | Monkeypox virus strain Yambuku_DRC_1985 | KP849471 | 197248 | 188 | 33.1 | NA | Democratic Republic of the Congo | 1985 | 13-May-15 | Viruses 7 (4), 2168-2184 (2015) |
| 65. | Monkeypox virus strain Zaire_1979-005 | DQ011155 | 196967 | 202 | 33.1 | Homo sapiens | Zaire, Democratic Republic of the Congo | NA (1979???) | 28-Sep-05 | J. Gen. Virol. 86 (PT 10), 2661-2672 (2005) |
| 66. | Monkeypox virus strain Zaire 1979-005 | HM172544 | 196959 | 197 | 33.1 | Homo sapiens | Zaire, Democratic Republic of the Congo | NA (1979???) | 25-Jul-10 | Virol. J. 7, 110 (2010) |
| 67. | Monkeypox virus Zaire-96-I-16 | NC 003310.1 | 196858 | 207 | 33.09 | NA | Zaire, Democratic Republic of the Congo | NA | 20-Dec-20 | FEBS Lett. 509 (1), 66-70 (2001) |
| 68. | Monkeypox virus MPX/UZ_REGA_1Belgium/2022 | ON622712 | 198010 | 211 | 32.9 | Homo sapiens | Belgium | 19-May-2022 | 22-May-2022 | Unpublished |
| 69. | Monkeypox virus MPX/UZ_REGA_2Belgium/2022 | ON622713 | 198016 | 211 | 32.9 | Homo sapiens | Belgium | 22-May-2022 | 22-May-2022 | Unpublished |
| 70. | Monkeypox virus MPXV_FRA_2022_TLS67 | ON602722.2 | 197103 | 219 | 33 | Homo sapiens | France | May-2022 | 25-May-2022 | Unpublished |
| 71. | Monkeypox virus MPXV_USA_2022_MA001 | ON563414 | 197205 | 211 | 33 | Homo sapiens | USA | May-2022 | 30-May-2022 | Unpublished |

### Genome Annotation

The genomes of 71 Monkeypox viruses were annotated using Prokka: rapid prokaryotic genome annotation [9]. To further refine the database, the prokka-genbank_to_fasta_db tool was used to generate the Monkeypox virus database. The genbank full format files of isolates namely Congo_2003_358, DRC_06-1070, MPXV-WRAIR7-61, COP-58, Israel_2018, Sudan_2005_01, Cote_d’Ivoire_1971, Liberia_1970_184, and Zaire-96-I-16 were downloaded from NCBI and the database was formatted. This database was further used to annotate the MPXV genomes in Prokka using --kingdom Viruses --gcode 1 flags.

### Phylogeny

The whole genome SNP-based phylogenetic tree was constructed using kSNP3 [10]. The optimum k-mer size of the dataset was determined using kchooser, a program that measures the diversity of sequences in the dataset. The whole genome consensus parsimony tree (-min_frac 0.5) was constructed in kSNP3 and visualized in iTOL [11]. Further, to access the similarity among the Monkeypox virus genomes, Average Nucleotide Identity was estimated using pyani [12]. The distance matrix was converted into a Neighbour Joining tree using the Ape package in R. Subsequently, the graph of the correlation matrix was plotted using the corrplot function in R. Additionally, to access the similarity based on core genome, the multifasta core genome alignment was generated using PRANK [13] in Roary [14] with minimum percentage identity of 95%. The alignment file was used to construct the phylogenetic tree using the maximum likelihood statistical method and Jukes-Cantor model in Mega XI [15].

### Core genome, pangenome and functional analysis

The GenBank files generated using prokka were subjected to the GET_HOMOLOGUES package for analysing the core genome using the OrthoMCL clustering algorithms with minimum identity and query coverage of 95% [16]. Furthermore, the substitution rates at non-synonymous and synonymous sites were determined for these core genomes. Using MUSCLE v3.8.31 and HyPhy v2.2.4 [17], orthologous gene clusters were aligned and stop codons were removed. For each orthologous gene cluster, Datamonkey v2.0 [18] (http://www.datamonkey.org/slac) used the single-likelihood ancestor counting (SLAC) method to calculate the dN/dS value. The dN/dS values were plotted using ggplot2 in R (R Development Core Team, 2015). Genes in monkeypox viruses were also analyzed for their differential presence. A gene absence matrix was generated using GenAPI [19]. Using ggplot2 in R, only genes that showed differential abundance were plotted. Furthermore, we clustered monkeypox proteins at 95% sequence similarity and query coverage using CD-HIT [20] and the resultant pangenome protein sequences were used for host-pathogen protein-protein interaction (PPI) analysis.

To access the functional potential, the genes were classified into six different categories viz. virulence and infection, genetic information processing, structural proteins, cell signalling and transduction, metabolism, and hypothetical proteins (Supplementary File 1). The presence of these genes was confirmed into six different categories across the different phylogenetic clades as determined by the whole genome SNP-based tree.

### Modeling of host-pathogen interaction network

Viruses rely on interactions between host and viral proteins to carry out all their life cycle functions, including infection, replication and even the assembly of new viral particles [7]. Monkeypox-virus protein sequences were submitted to HPIDB3.0 [21], the host-pathogen interaction database, to predict their direct interaction with humans as the principal host. BLASTp was used to retrieve homologous host/pathogen protein sequences [22]. For high-throughput analysis, BLASTp was used to search multiple protein sequences at once, and the results were presented both in a tabular format and as sequence alignments [21, 22]. Cytoscape v3.9.1 was used to construct and visualize the HPI network [23]. As the constructed network demonstrated, proteins with the highest degrees that interact with a large number of signalling proteins, played a key regulatory role as hubs. The hub proteins were identified using Network Analyzer [24], a plugin of Cytoscape v3.9.1.

### Computational structure analysis of key monkeypox viral protein

For understanding the effects of mutations on the stability of key viral proteins, computational structural analysis was performed on the one key viral protein in the PPI network cytokine response-modifying protein B (CrmB). The computational structure of wild-type and mutant CrmB were constructed using the Phypre2 [25] and Swiss model [26]. The structure was energy minimized by the Chiron energy minimization server [27]. The structure was also validated using the Ramachandran plot. The effect of the mutation was analyzed using FoldX [28]. The structures were repaired before building the mutant models. Repair protein structure helps identify those residues with poor torsion angles, or VanderWaal’s classes, or total energy, and repairs them. The FoldX tool provides the difference in Gibbs energy of the protein.

## Results

### General Genomic Attributes

The monkeypox virus belongs to genus Orthopoxvirus with the average genome size and coding sequences of 196.44 ± 3.42 Kb and 188.57 ± 10.43, respectively, whereas the average %GC was 33.06 ± 0.06% (Table 1). The results are in consensus with previously published reports [29]. The majority of isolates studied so far have been isolated from Central & West African countries, major ones from the Democratic Republic of Congo (formerly known as Zaire), Israel, the USA, Singapore and France (Table 1). In all likelihood, the disease was transmitted via exotic animals transported from tropical rainforests to other parts of the western hemisphere [30]. Interestingly, we also examined four isolates from the current outbreak (MPX/UZ_REGA_1Belgium/2022, MPX/UZ_REGA_2Belgium/2022, MPXV_FRA_2022_TLS67 and MPXV_USA_2022_MA001). Two of these isolates were reported from Belgium, one each from France and the USA (Table 1). The largest genome size was observed in the case of strain Sudan 2005_01 (206.37 kb), whereas the smallest was in case of strain MPXV-M5320 M15 Bayelsa (185.31 kb), which is in agreement with the previous reports [29]. Interestingly, the highest number of coding sequences/ORFs (n=219) were seen in case of strain MPXV_FRA_2022_TLS67 with a genome size of 197.12kb. Out of the 71 isolates, 52 were isolated from *Homo sapiens* between the years 2005 and 2022, while one each was from a wild monkey, dormouse, Gambian rat, rope squirrel and Prairie Dog (Table 1), suggesting the potential of monkeypox virus to infect a wide array of hosts.

### Phylogenetic Analysis

A phylogenetic tree based on whole genome SNPs representing 16 different viruses was constructed for the members of the genus Orthopoxvirus, including Ectromelia (n=3), Orthopoxvirus Abatino (n=1), Akhmeta (n=4), Cetacean (n=1), Raccoonpox (n=1), Skunkpox (n=1), Volepox (n=1), Cowpox (n=35), Variola (n=4), Tetrapox (n=1), Camelopox (n=9), Vaccinia (n=15), Horsepox (n=2), Rabbitpox (n=1), Buffalopox (n=6) and Monkeypox (n=71) viruses. The Monkeypox virus clustered distinctly from other members of the genus *Orthopoxvirus* but closely with the Vaccinia virus (Figure 1A). The Vaccinia virus clade comprised of other viruses namely Horsepox virus, Rabbitpox virus and Buffalopox virus.

**Figure 1:**
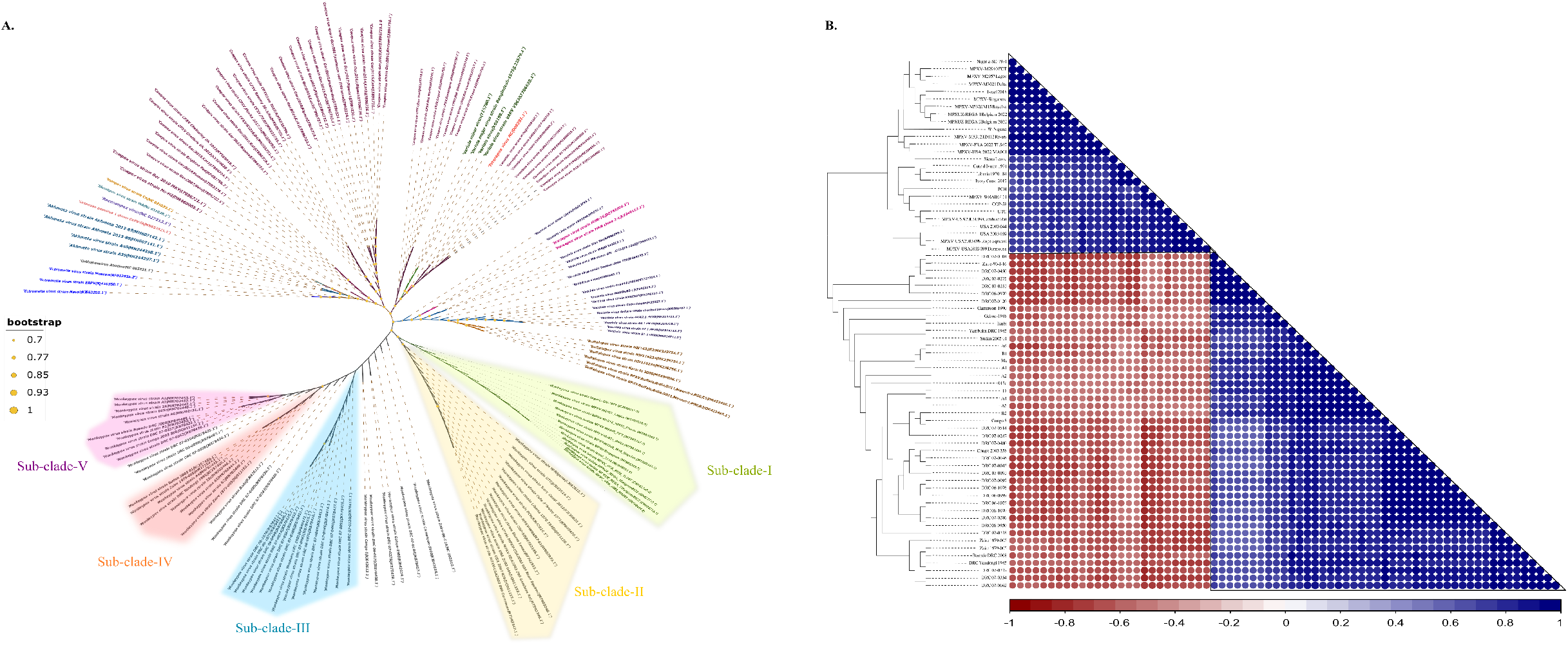
The Phylogenetic analysis of monkeypox virus A. The whole genome SNP based phylogenetic tree of members of Orthopox virus was constructed using kSNP3 and visualized in iTOL. B. The average nucleotide identity matrix representing the sequence similarity of 71 monkeypox viruses. The distance matrix was converted into Neighbour Joining tree using the Ape package in R. Subsequently, the graph of correlation matrix was plotted using the corrplot function in R.

There were similar clustering patterns in the monkeypox virus Clade I in the whole genome-based SNP tree, ANI-based dendrogram (Figure 1B) and core genome tree (Figure 2A) except isolate W-Nigeria (KJ642615.1), which clustered with clade II members in whole genome SNP tree, suggesting at the genome level this isolate could have accumulated more variations. Most of the isolates in clade I originated from Nigeria. Interestingly, two isolates that had been reported from Singapore (MT9033421) and Israel (MN648051.1), also clustered with the Nigerian isolates, suggesting the high level of genomic similarity between these two isolates and the Nigerian isolates. A striking finding was the clustering of all the newly reported isolates together from 2022 with those in clade I (Figure 1A) in close proximity with strain Israel 2018 (MN 648051.1), suggesting that they emerged from those in clade I. However, further reports will clarify the pattern of emergence of these isolates.

**Figure 2:**
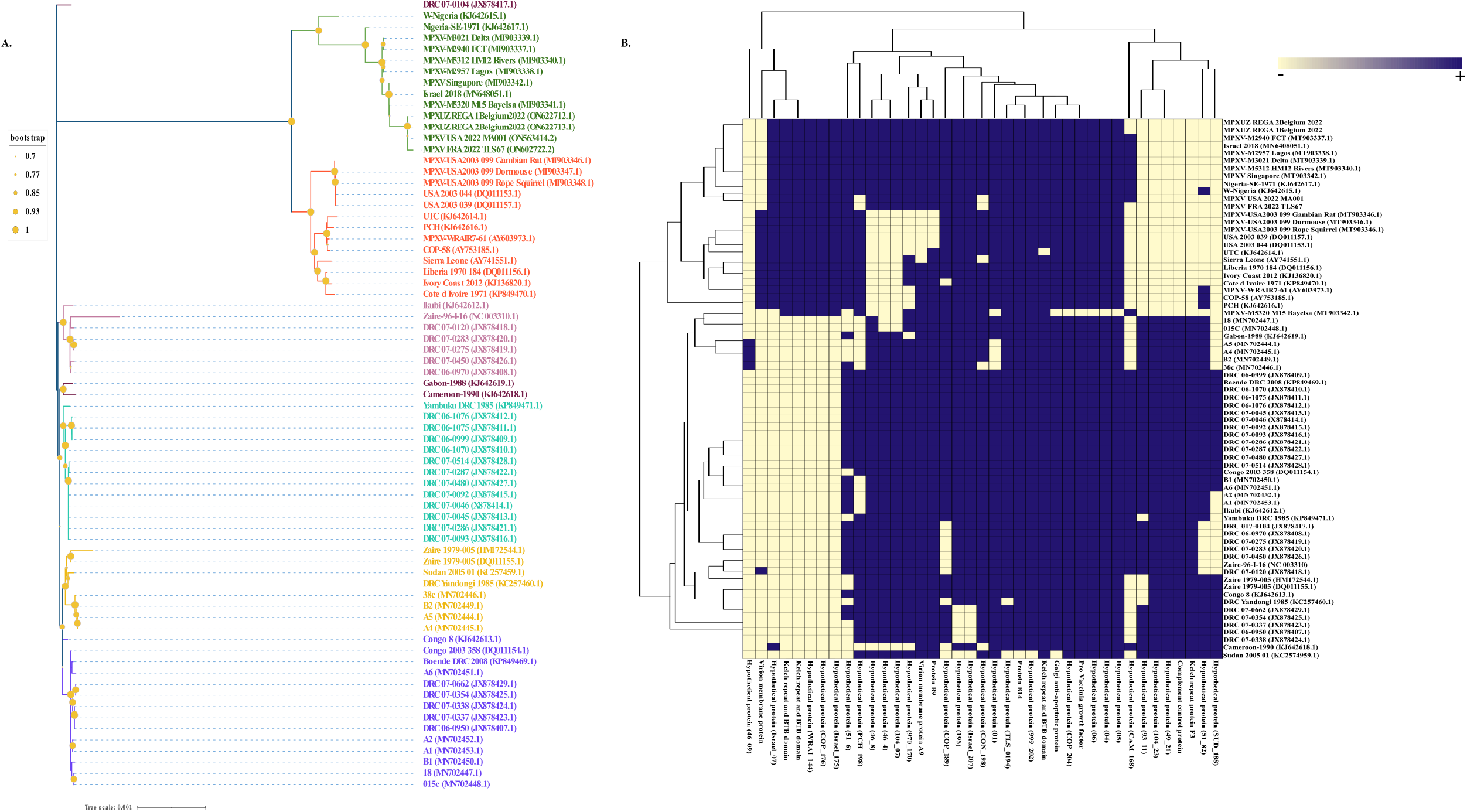
A. The phylogenetic tree constructed using the core genome (n=140). The multifasta core genome alignment was generated using PRANK in Roary with minimum percentage identity of 95%. The alignment file was used to construct the phylogenetic tree using Maximum Likelihood statistical method and Jukes-Cantor model in Mega XI.

Clade II, on the other hand, shows the consensus tree topology across all three methods used for the delineation of these species. This clade, however, harbours monkeypox virus from a variety of hosts, such as wild monkeys (Ivory Coast 2012 (KJ136820)), rope squirrels (USA2003_099_Rope_Squirrels (MT903348)), Gambian rats (USA2003_099_Gambian_Rats (MT903346)), dormouse (USA2003_099_Dormouses (MT903347)), prairie dog (USA_2003_044 (DQ011153)). The latter four isolates were found in the United States. Interestingly, one of the strains USA_2003_039 (DQ011157) isolated from *Homo sapiens* clusters tightly with different host types, namely rope squirrels, Gambian rats, dormouse, and prairie dogs, implying a recent emergence in humans and the infection is acquired through contact with animals. Additionally, the isolates from the Netherlands (strain UTC (KJ642614)) and France (strain PCH (KJ642616)) clustered separately based on the whole genome SNP method, but clustered together in core genome (Figure 2A) and ANI-based phylogeny (Figure 1B), suggesting the accumulation of additional mutations in due course.

Further, the Clade III in whole genome SNP-based tree clustered monkeypox viruses originated from the Democratic Republic of Congo and the limited metadata suggests that the majority of these isolates are of human origin. But the consensus tree topology was observed only in the case of phylogenetic trees derived using whole-genome SNPs and core genomes, except for strain DRC-07-120 (JX878418) which formed a minor clade with strain Zaire-96-I-16 (DQ011155). In both WGS, SNP-based and core genome trees, Clade IV clustered the isolates from Zaire, Democratic Republic of Congo, and the Central African Republic, all of which were of human origin, indicating that the infection spread by traveling and human contact. Like Clade IV, Clade V clustered the strains from the Democratic Republic of Congo and the Central African Republic and showed consensus tree topology in WGS SNP-based and core genome trees.

Additionally, the most ancestral strain in the core genome phylogenetic tree was the Monkeypox virus strain DRC 07-0104 (JX878417), while the strain Nigeria-SE-1971 (KJ642617) exhibited ancestry with Vaccinia virus.

### Functional and evolutionary dynamics of monkeypox virus

First, we examined the genes which were differentially enriched in 71 isolates of monkeypox. A total of 39 genes showed differential abundance (Figure 2B). The fact that 28 of these 39 genes code for hypothetical proteins and on an average 123 genes code for hypothetical proteins per genome indicates that experimental evidence is required to ascertain their role in the viral genome. Further, the genes which were differentially enriched include protein B9, virion membrane protein, protein B14, Golgi-antiapoptotic protein, Kelch repeat protein F3, complement control protein and Kelch repeat and BTB domain (Figure 2B). Multiple reports have suggested that poxviruses encode for kelch/BTB proteins (cell signaling and transduction group) and these domains aid in substrate recruitment to cullin-3-based ubiquitin ligases. These are a part of the survival strategies of the virus against the host’s antiviral responses and immune evasion [31]. Complement control proteins (CCP) are also conserved in most poxviruses and genomic comparisons of viral isolates from Congo and West Africa have revealed that increased virulence in some isolates is credited to these groups of proteins. CCP homologs have been identified in various orthopox viruses such as vaccinia, variola, ectromelia, and cowpox virus. They have shown to be capable of inhibiting the activity of the complement system via binding of proteins and speeding up the decay of the various convertase enzymes in both classical and alternative pathways [32]. Golgi anti-apoptotic proteins (GAAP), first discovered in the camelpox virus are hydrophobic proteins of the Golgi membranes. Their function is to protect the cell from apoptotic stimuli and regulation of Ca^2+^ fluxes. It is postulated that GAAPs have a role in Ca^2+^ signaling and anti-apoptotic activities. They are a part of the transmembrane Bax inhibitor-containing motif (TMBIM) family that carries out similar functions [33]. As seen in Figure 2B, the maximum MPX under study has copies of GAAPs. Further, these proteins differ at a high sequence similarity level (>95%) and gene variants with low similarity may be present in these isolates, which would provide a novel avenue for further research on monkeypox viruses.

Second, we analysed the core genome of Monkeypox virus. A total of 140 genes were conserved among the monkeypox viruses (Figure 3A, Supplementary Table 2). Among the 140 conserved genes, 34 code for hypothetical proteins. We then specifically used the core genome to quantifying selection pressure. The dN/dS metric is one of the most widely used methods of quantifying selection pressure which compares synonymous and non-synonymous substitutions. This method is frequently used to determine whether the protein is subject to purifying selection (dN/dS <1), evolving neutrally (dN/dS ≈ 1), or undergoing positive, diversifying selection (dN/dS >1)[34]. In the monkeypox core genome, we identified five proteins that were showing diversifying selection. These proteins include K3L, F1, CrmB, envelop protein and putative FAD-linked protein (Figure 3B). The K3L protein shares 28% homology with eukaryotic translation initiation factor 2 (eIF2α) which is a substrate for Protein Kinase R (PKR)[35]. As a component of the innate immune system in vertebrates, the PKR when interacts with K3L (pseudo-substrate inhibitor), magnifies the conundrum posed by viral mimicry. The PKR’s effectiveness depends on its interaction with eIF2α rather than it mimics such as K3L. K3L (dN/dS ≈ 1.13) is still working towards achieving the optimal state of mimicry, which could help the virus to evade the innate immune response. The result agrees with the previous report by Elde *et al* (2009)[36], which also reported the unchanged nature of eIF2α in simian primates and further suggested the evolution of K3L in response to the adaptive changes in PKR. Additionally, the mitochondrial-associated inhibitor of apoptosis i.e., protein F1 is positively selected and it has been reported to block apoptosis by binding to Bak in the vaccinia virus [37]. Further, it has been demonstrated that the deletion of the F1L gene from the vaccinia genome increased apoptosis during infection [37] and promotes virulence by inhibiting inflammasome activation [38], and its greater purifying selection (dN/dS ≈ 1.61) indicates that the gene is evolving in monkeypox virus during evolutionary processes in order to enhance its virulence. Likewise, the CrmB protein (dN/dS ≈ 1.61) is also under purifying selection, which may provide a selective advantage to the monkeypox virus to protect the infected cells from the host antiviral response. A cytokine secreted by T cells and macrophages, TNF-α protects cells from viral infection and can kill infected cells [39]. The CrmB protein binds to TNF-α and TNF-β and thus protects the infected cells by preventing TNF-mediated immune response against viruses [39]. Furthermore, the two other positively selected proteins, i.e., the envelope protein and the putative FAD-linked protein, might be evolving to help the virus to overcome intracellular host restrictions and achieve efficient survival.

**Figure 3:**
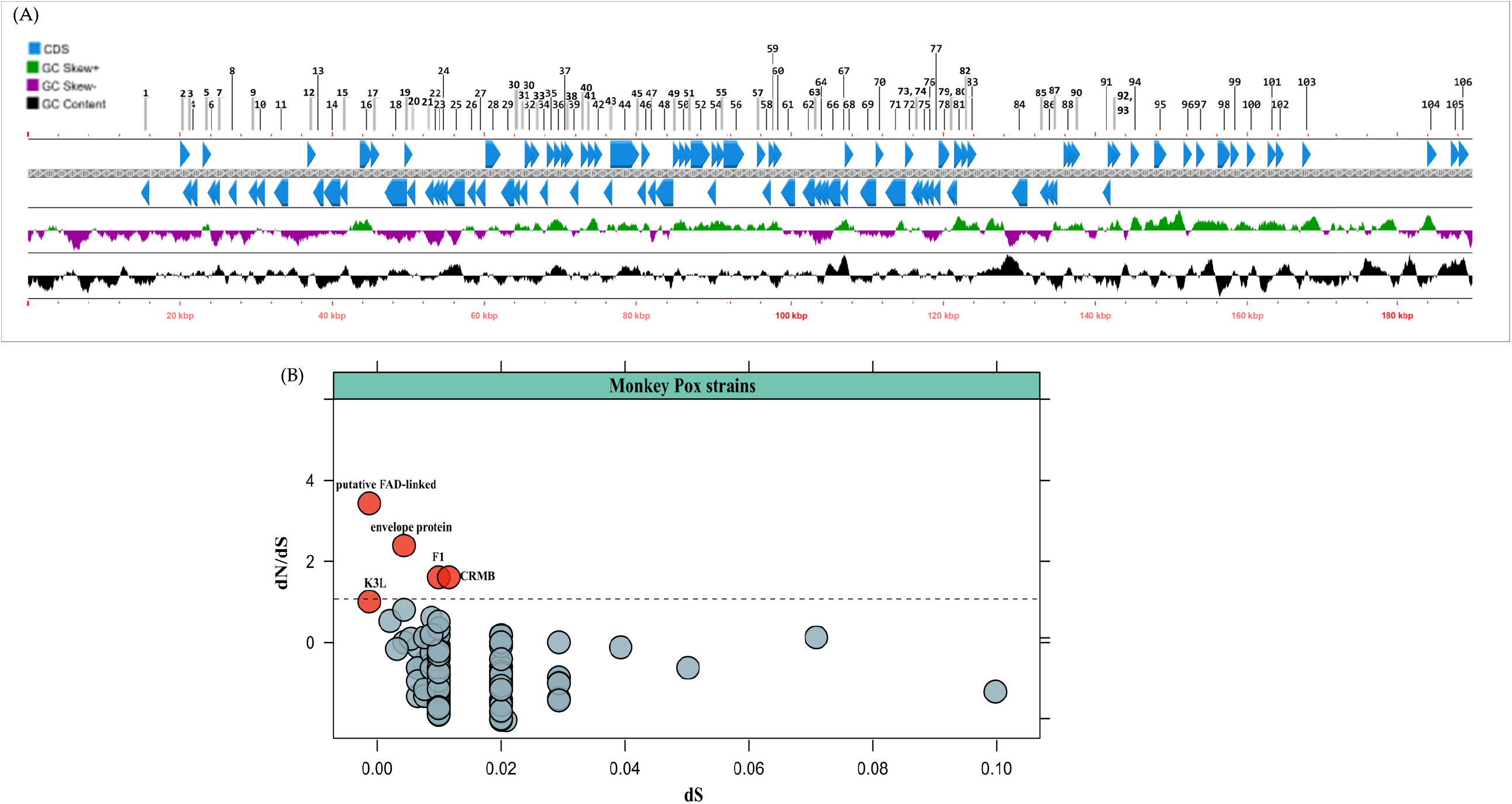
A. The mapping of core gene distribution across the reference genome of Monkeypox virus strain 015. B. The dN/dS analysis of core genes for estimating the direction of selection.

As mentioned above, MPXV genes were broadly divided into five broad categories for functional analysis (Figure 4, Supplementary Table 1). The maximum number of genes were characterised under the category virulence and infection (44.46± 1.39), followed by genetic information processing (41.80 ± 0.87), structural proteins (23.07± 0.97), cell signalling and transduction (20.30 ± 1.29) while the lowest were involved in metabolism (16.3 ± 0.57), emphasizing the dependency of viral genomes on host profiles for metabolism processing, while converging more on conferring virulence. There were differences in the number of genes under different categories at the isolate level, but no significant differences were observed at the clade level (Figure 4). Interestingly, the mutation analysis in clade I revealed 59 proteins with amino acid level mutations (Supplementary Table 3), suggesting the proteins are constantly adapting to their broad host range.

**Figure 4:**
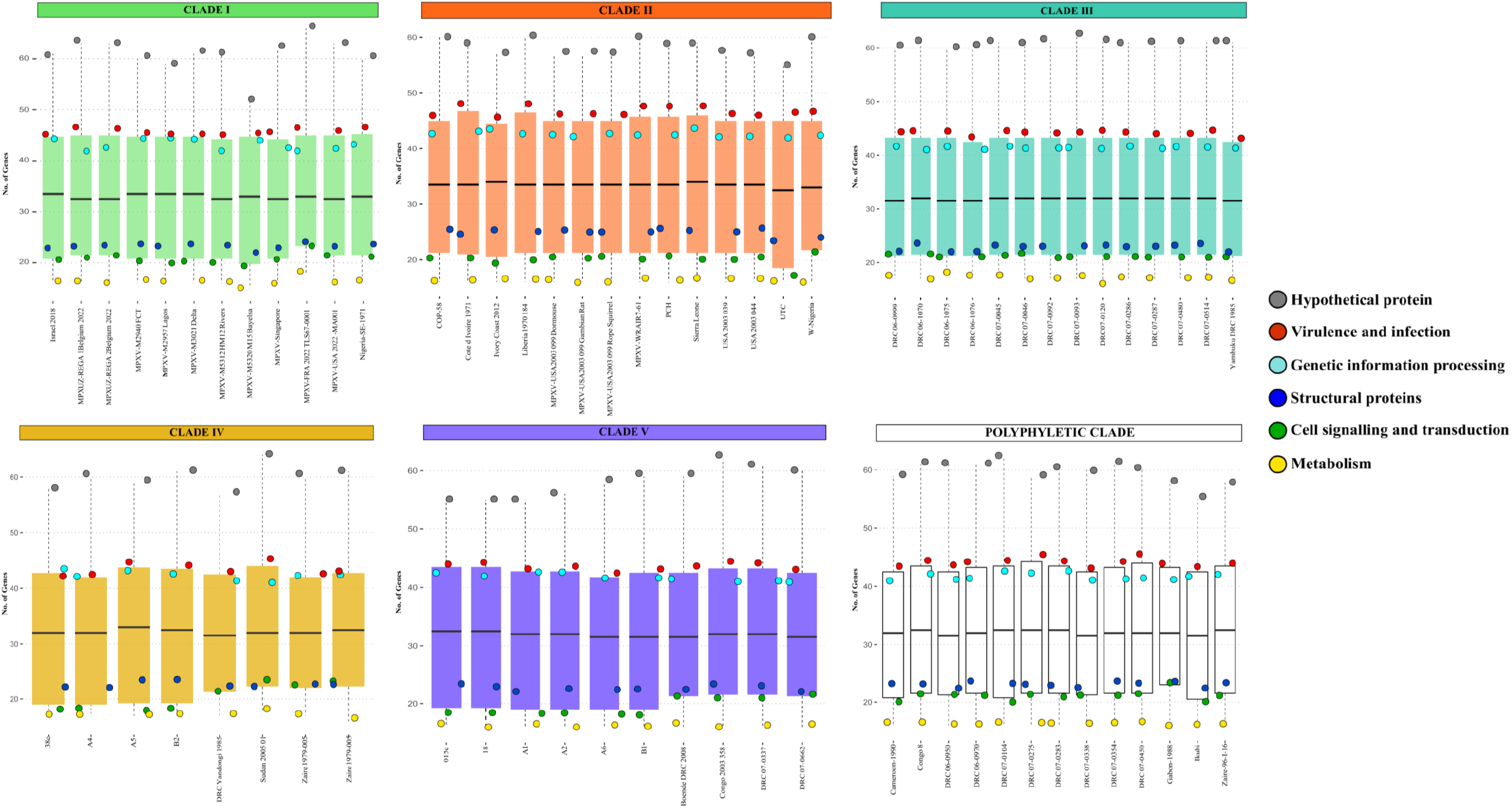
The distribution of monkeypox proteins in different phylogenetic groups under six broad categories namely Genetic information processing, Cell signalling & transduction, Metabolism, Virulence and infection and Structural proteins.

### Exploring the network for host-pathogen interaction between monkeypox virus and its host

A detailed study of the interplay between poxvirus proteins and the host immune system is of great interest to understand the infectivity of these viruses and will also pave the way to successfully design and administer recombinant vaccines. The HPI network of Monkeypox-virus contained 331 edges and 273 nodes, including 55 viral and 218 host proteins (Figure 5A). The significant existence of a few hubs, namely, protein E3 and serine protease inhibitor-2 (SPI-2), protein K7, and CrmB in the network and the attraction of a large number of low-degree nodes toward each hub showed strong evidence of control of the topological properties of the network by a few hub proteins. Protein E3 was found to have a connection with 47 human host proteins whereas SPI-2, protein K7, and CrmB exhibited 37, 33, and 21 degrees, respectively (Figure 5A). These monkeypox viral proteins were the main hubs in the network, which regulate/control the network. Based on degree distribution, the viral protein E3 showed the highest interaction, followed by Serine protease inhibitor 2, protein K7, and CrmB.

**Figure 5:**
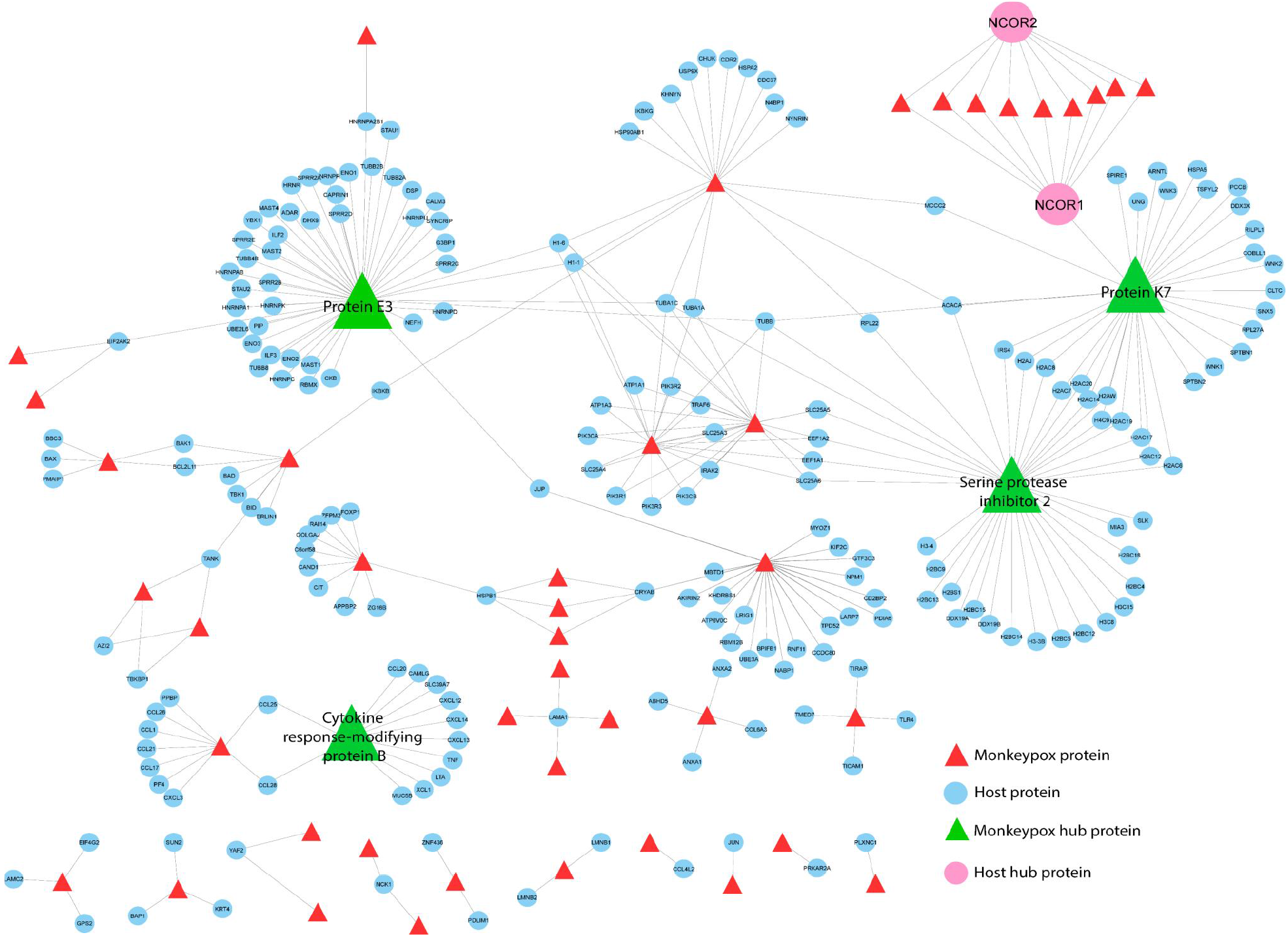
The host-pathogen interaction network. Monkeypox-virus protein sequences were submitted to HPIDB3.0, the host-pathogen interaction database, to predict their direct interaction with humans as the principal host. Cytoscape v3.9.1 was used to construct and visualize the HPI network. The hub proteins were identified using Network Analyzer, a plugin of Cytoscape v3.9.1.

Protein E3 plays a critical role in unhindered viral replication by blocking the cellular innate immune system [40]. Generation of interferons (IFN) is the prime response against viral infection. E3 protein of monkeypox virus is known to produce IFN resistant phenotypes by inhibiting the phosphorylation of PKR and eIF2α [41] which is in accordance with our interactome study where E3 is interacting with host EIF2AK2. Further, the interactome analysis also showed that E3 protein interacts with host interleukin enhancing binding factors 2 and 3 (ILF2/3) which are known for providing an innate antiviral response by regulating the transcription of IL2 gene during T cell activation [42]. Furthermore, it is known that the suppression of cognate T cell activation which evades CD8+ and CD4+ responses is one of the immune escape mechanisms of monkeypox virus [43].

Our study has shown that viral protein K7 is interacting with several host proteins like NCOR1, SPIRE1, DDX3X, WNK1/2/3, SNX5, etc. K7 is known as an antagonist of innate immunity and acts as a virulence factor that inhibits IRF3 and NFKB activation [44]. Studies on the vaccinia virus have also shown that K7 protein interacts with SPIR-1 which is a virus restriction factor and activates innate immune signaling which is critical for the host response against viral infection [45].In addition to it, our interactome analysis showed the interaction between K7 and DEAD-box protein 3 (DDX3). In the viral infection, the viruses are detected by several pattern recognition receptors (PRR) like TLRs, RIG like helicases, etc., which promotes antiviral activity by inducing IFN-production through the activation of interferon (IFN)-regulatory factor 3 (IRF3) and IRF7. Studies on vaccinia virus (VACV) have shown that interaction between K7 protein with DDX3 inhibits PRR-induced IFN-β induction by suppressing TBK1/IKKε-mediated IRF activation [46].

K7 of vaccinia virus also interacts with WNK (with-no-lysine) family which plays a crucial role in antiviral immune response and knockdown of WNK family members resulting in increased growth of vaccinia virus in the host. WNK 1 and WNK 3 stimulate interleukin 1(IL-1) by activating p38 kinase which is inhibited by co-expression of K7 [47]. Interestingly, our interactome study between human and monkeypox virus also showed an interaction between K7 and WNK family members. The PPI interactions also showed that two host proteins, NCOR1 and NCOR2, exhibit a maximum interaction with viral proteins. NCOR1 interacting with viral hub protein K7 showed an important co-relation with infections caused by the monkeypox virus. Both NCOR1 and NCOR2 are responsible for the repression of transcription by promoting histone deacetylation and chromatin repression and impeding access to transcription factors [48] and hence can be proposed to bring about gene silencing during infection by the viruses in the host.

In the activation of antiviral immune response by the host, TNF plays a pivotal role, where it either acts directly as cell death-inducing cytokine on virus-infected cells or indirectly as an inducer of the innate and adaptive immune response against the invading virus. Viruses have co-evolved with the host for their better survival and devised several strategies to evade TNF-mediated responses. One such strategy is the generation of Cytokine response modifying protein B (CrmB) which is an excellent immunomodulator. Studies have shown that CrmB binds with TNF and several chemokines like CCL25, CCL28, CXCL12β, CXCL13 and CXCL14 to inhibit host immune responses against the virus [48, 49]. The binding of CrmB with chemokines prevents recruitment of T cells and B cells, dendritic cell migration to epidermal tissue and recruitment of B cells to spleen and lymph nodes [50]. Interestingly, our PPI results also showed the interaction of CrmB protein of monkeypox virus with host TNF and several chemokines like CCL1, CCL21, CCL25, CXCL13, CXCL14, etc. Another key protein hub, SPI-2 also contributes to poxvirus immune escape. By targeting caspase-1, SPI-2 prevents apoptosis and cytokine activation. Further, the induction of IFN-β and its downstream genes is inhibited by the ectopic expression of SPI-2, thus preventing the host to confer INF mediated immune response against viruses.

### Structural analysis of CrmB protein

Computational structural analysis has been carried out to understand the effect of mutation on the key CrmB viral proteins. Based on the host-pathogen analysis, a few key proteins like protein E3, SPI-2, protein E8, and CrmB seemed to generate interest. Among these key proteins, the structural analysis of CrmB was performed, as this gene was showing positive selection in dN/dS analysis. In the case of CrmB, a structurally similar template was not identified for the major part of the protein. So, modeling was carried out using Phyre2 tool to obtain a 3D structure of the CrmB protein (Figure 6). The hydrogen bond analysis using the pymol showed that due to the insertion of valine and S54F mutation there is a loss of some hydrogen bond which could disturb the structure (Figure 6A). The stability (ΔG) of a protein is defined by the free energy, which is expressed in kcal/mol. The lower the value of ΔG, the more stability. Here we calculated the ΔΔG which is the difference in free energy between wild-type and mutant. The mutation that brings energy (ΔΔG) higher than 0 will destabilize the structure and lower than 0 will stabilize the structure. In the case of CrmB protein, the four mutations were S121A, S54F, R224V and an insertion of valine (V) at position 172. Among all the four mutations, three of them S121A, S54F and insertion of V at 172 were destabilizing the structure, whereas R224V was found to stabilize the protein (Figure 6B).

**Figure 6:**
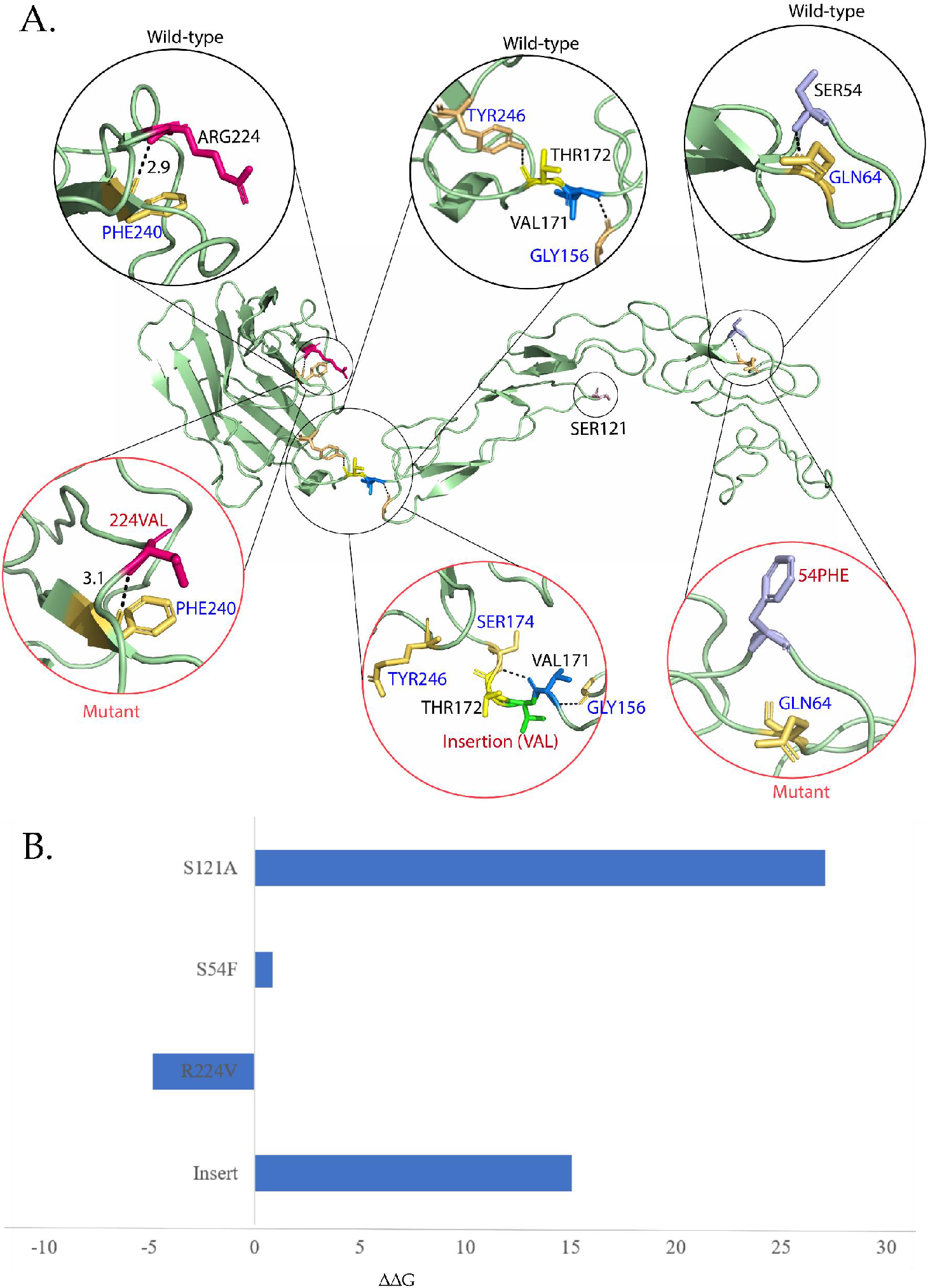
The structure analysis of crmB protein. A) Structures depicting the mutations i.e., S121A, S54F, R224V and an insertion of valine (V) at position 172; B) The ΔΔG analysis to depict the stability of mutations. Mutations that bring energy (ΔΔG) higher than 0 will destabilize the structure and mutations that bring energy (ΔΔG) lower than 0 will stabilize it.

CrmB protein has a Smallpox virus-encoded chemokine receptor (SECRET) domain that dispenses chemokine inhibitory activities and that allows the virus to differentially block chemokines and TNFs [50]. CrmB host protein interacts with the host TNFs via its N-terminal domain [51]. It also interacts with Calcium Modulating Ligand (CAMLG) which takes part in the calcium signal transduction pathway [50]. Further, it interacts with Mucin 5B protein, a glycosylated macromolecular component of mucus secretions. The protein K7, a part of the virulence and infection proteins interacts with the DEAD-box RNA helicase DDX3, tumour necrosis factor-associated factor 6 and interleukin-1 receptor associated kinase to inhibit activity of interferon regulatory factors [44].

## Conclusion

With the sudden increase in the number of cases of human monkeypox all over the world, it is necessary that we rapidly unfold the mutational rate, virulence and evolutionary lineage of different isolates from across different geographical locations of the world from where cases have been reported. Although we have only analyzed 71 genomes but one of the major outcomes of this study was the fact that the four strains associated with the recent outbreak were not just clustered in Clade I which was predominantly occupied by Nigerian isolates. On the contrary these strains exhibited maximum similarity with an Israelian isolate. Another interesting outcome sprouted from the Host-Pathogen interactome analysis, common CrmB protein presumptively appeared to be a key regulator in conferring monkeypox virus with a selective advantage against the host immune system. The presence of S54F mutation in CrmB protein which was common of all the recently documented isolates of monkeypox, reflected that the selected strains are possibly adapting. The current outbreak may end up bringing another pandemic, this it is imperative that emphasis should be shifted to sequencing more genomes of monkeypox virus from across different geographical locations to devise better treatment and prevention strategies and to prevent another major outbreak.

## ACKNOWLEDGMENTS

RK acknowledges Magadh University, Bodh Gaya, for providing support. RL and US also acknowledge The National Academy of Sciences, India, for support under the NASI-Senior Scientist Platinum Jubilee Fellowship Scheme.

## Author contribution statement

RK, IKS and RL conceived and designed the study. RK, SN, SH, US, KP, GGD, SA, AD, MS executed the analysis and prepared figures. RK, SN, SH, US, KP, GGD, SA, AD, MS, MS, IKS and RL wrote the manuscript and finalized the drafts.

## REFERENCES

1. Ladnyj ID, Ziegler P, Kima E: A human infection caused by monkeypox virus in Basankusu Territory, Democratic Republic of the Congo. Bull World Health Organ 1972, 46(5):593–597.

2. Sklenovska N, Van Ranst M: Emergence of Monkeypox as the Most Important Orthopoxvirus Infection in Humans. Front Public Health 2018, 6:241.

3. Adler H, Gould S, Hine P, Snell LB, Wong W, Houlihan CF, Osborne JC, Rampling T, Beadsworth MB, Duncan CJ et al: Clinical features and management of human monkeypox: a retrospective observational study in the UK. Lancet Infect Dis 2022.

4. Vaughan A, Aarons E, Astbury J, Brooks T, Chand M, Flegg P, Hardman A, Harper N, Jarvis R, Mawdsley S et al: Human-to-Human Transmission of Monkeypox Virus, United Kingdom, October 2018. Emerg Infect Dis 2020, 26(4):782–785.

5. Reynolds MG, Yorita KL, Kuehnert MJ, Davidson WB, Huhn GD, Holman RC, Damon IK: Clinical manifestations of human monkeypox influenced by route of infection. J Infect Dis 2006, 194(6):773–780.

6. Kumar R, Verma H, Singhvi N, Sood U, Gupta V, Singh M, Kumari R, Hira P, Nagar S, Talwar C et al: Comparative Genomic Analysis of Rapidly Evolving SARSCoV-2 Reveals Mosaic Pattern of Phylogeographical Distribution. mSystems 2020, 5(4).

7. Gupta V, Haider S, Verma M, Singhvi N, Ponnusamy K, Malik MZ, Verma H, Kumar R, Sood U, Hira P et al: Comparative Genomics and Integrated Network Approach Unveiled Undirected Phylogeny Patterns, Co-mutational Hot Spots, Functional Cross Talk, and Regulatory Interactions in SARS-CoV-2. mSystems 2021, 6(1).

8. Gurevich A, Saveliev V, Vyahhi N, Tesler G: QUAST: quality assessment tool for genome assemblies. Bioinformatics 2013, 29(8):1072–1075.

9. Seemann T: Prokka: rapid prokaryotic genome annotation. Bioinformatics 2014, 30(14):2068–2069.

10. Gardner SN, Slezak T, Hall BG: kSNP3.0: SNP detection and phylogenetic analysis of genomes without genome alignment or reference genome. Bioinformatics 2015, 31(17):2877–2878.

11. Letunic I, Bork P: Interactive Tree Of Life (iTOL) v5: an online tool for phylogenetic tree display and annotation. Nucleic Acids Res 2021, 49(W1):W293–W296.

12. Pritchard L, Glover RH, Humphris S, Elphinstone JG, Toth IK: Genomics and taxonomy in diagnostics for food security: soft-rotting enterobacterial plant pathogens. Analytical Methods 2016, 8(1):12–24.

13. Loytynoja A: Phylogeny-aware alignment with PRANK. Methods Mol Biol 2014, 1079:155–170.

14. Page AJ, Cummins CA, Hunt M, Wong VK, Reuter S, Holden MT, Fookes M, Falush D, Keane JA, Parkhill J: Roary: rapid large-scale prokaryote pan genome analysis. Bioinformatics 2015, 31(22):3691–3693.

15. Tamura K, Stecher G, Kumar S: MEGA11: Molecular Evolutionary Genetics Analysis Version 11. Molecular Biology and Evolution 2021, 38(7):3022–3027.

16. Contreras-Moreira B, Vinuesa P: GET_HOMOLOGUES, a versatile software package for scalable and robust microbial pangenome analysis. Appl Environ Microbiol 2013, 79(24):7696–7701.

17. Pond SL, Frost SD, Muse SV: HyPhy: hypothesis testing using phylogenies. Bioinformatics 2005, 21(5):676–679.

18. Weaver S, Shank SD, Spielman SJ, Li M, Muse SV, Kosakovsky Pond SL: Datamonkey 2.0: A Modern Web Application for Characterizing Selective and Other Evolutionary Processes. Mol Biol Evol 2018, 35(3):773–777.

19. Gabrielaite M, Marvig RL: GenAPI: a tool for gene absence-presence identification in fragmented bacterial genome sequences. BMC Bioinformatics 2020, 21(1):320.

20. Li W, Godzik A: Cd-hit: a fast program for clustering and comparing large sets of protein or nucleotide sequences. Bioinformatics 2006, 22(13):1658–1659.

21. Ammari MG, Gresham CR, McCarthy FM, Nanduri B: HPIDB 2.0: a curated database for host-pathogen interactions. Database (Oxford) 2016, 2016.

22. Altschul SF, Gish W, Miller W, Myers EW, Lipman DJ: Basic local alignment search tool. J Mol Biol 1990, 215(3):403–410.

23. Shannon P, Markiel A, Ozier O, Baliga NS, Wang JT, Ramage D, Amin N, Schwikowski B, Ideker T: Cytoscape: a software environment for integrated models of biomolecular interaction networks. Genome Res 2003, 13(11):2498–2504.

24. Assenov Y, Ramirez F, Schelhorn SE, Lengauer T, Albrecht M: Computing topological parameters of biological networks. Bioinformatics 2008, 24(2):282–284.

25. Kelley LA, Mezulis S, Yates CM, Wass MN, Sternberg MJE: The Phyre2 web portal for protein modeling, prediction and analysis. Nat Protoc 2015, 10(6):845–858.

26. Waterhouse A, Bertoni M, Bienert S, Studer G, Tauriello G, Gumienny R, Heer FT, de Beer TA P, Rempfer C, Bordoli L et al: SWISS-MODEL: homology modelling of protein structures and complexes. Nucleic Acids Research 2018, 46(W1):W296–W303.

27. Ramachandran S, Kota P, Ding F, Dokholyan NV: Automated minimization of steric clashes in protein structures. Proteins 2011, 79(1):261–270.

28. Buß O, Rudat J, Ochsenreither K: FoldX as Protein Engineering Tool: Better Than Random Based Approaches? Computational and Structural Biotechnology Journal 2018, 16:25–33.

29. Kugelman JR, Johnston SC, Mulembakani PM, Kisalu N, Lee MS, Koroleva G, McCarthy SE, Gestole MC, Wolfe ND, Fair JN et al: Genomic variability of monkeypox virus among humans, Democratic Republic of the Congo. Emerg Infect Dis 2014, 20(2):232–239.

30. Ligon BL: Monkeypox: a review of the history and emergence in the Western hemisphere. Semin Pediatr Infect Dis 2004, 15(4):280–287.

31. Wilton BA, Campbell S, Van Buuren N, Garneau R, Furukawa M, Xiong Y, Barry M: Ectromelia virus BTB/kelch proteins, EVM150 and EVM167, interact with cullin-3-based ubiquitin ligases. Virology 2008, 374(1):82–99.

32. Hudson PN, Self J, Weiss S, Braden Z, Xiao Y, Girgis NM, Emerson G, Hughes C, Sammons SA, Isaacs SN et al: Elucidating the role of the complement control protein in monkeypox pathogenicity. PLoS One 2012, 7(4):e35086.

33. Saraiva N, Prole DL, Carrara G, Maluquer de Motes C, Johnson BF, Byrne B, Taylor CW, Smith GL: Human and viral Golgi anti-apoptotic proteins (GAAPs) oligomerize via different mechanisms and monomeric GAAP inhibits apoptosis and modulates calcium. J Biol Chem 2013, 288(18):13057–13067.

34. Verma H, Kumar R, Oldach P, Sangwan N, Khurana JP, Gilbert JA, Lal R: Comparative genomic analysis of nine Sphingobium strains: insights into their evolution and hexachlorocyclohexane (HCH) degradation pathways. BMC Genomics 2014, 15:1014.

35. Kawagishi-Kobayashi M, Silverman JB, Ung TL, Dever TE: Regulation of the protein kinase PKR by the vaccinia virus pseudosubstrate inhibitor K3L is dependent on residues conserved between the K3L protein and the PKR substrate eIF2alpha. Mol Cell Biol 1997, 17(7):4146–4158.

36. Elde NC, Child SJ, Geballe AP, Malik HS: Protein kinase R reveals an evolutionary model for defeating viral mimicry. Nature 2009, 457(7228):485–489.

37. Postigo A, Cross JR, Downward J, Way M: Interaction of F1L with the BH3 domain of Bak is responsible for inhibiting vaccinia-induced apoptosis. Cell Death Differ 2006, 13(10):1651–1662.

38. Gerlic M, Faustin B, Postigo A, Yu EC, Proell M, Gombosuren N, Krajewska M, Flynn R, Croft M, Way M et al: Vaccinia virus F1L protein promotes virulence by inhibiting inflammasome activation. Proc Natl Acad Sci U S A 2013, 110(19):7808–7813.

39. Weaver JR, Isaacs SN: Monkeypox virus and insights into its immunomodulatory proteins. Immunol Rev 2008, 225:96–113.

40. White SD, Jacobs BL: The amino terminus of the vaccinia virus E3 protein is necessary to inhibit the interferon response. J Virol 2012, 86(10):5895–5904.

41. Arndt WD, Cotsmire S, Trainor K, Harrington H, Hauns K, Kibler KV, Huynh TP, Jacobs BL: Evasion of the Innate Immune Type I Interferon System by Monkeypox Virus. J Virol 2015, 89(20):10489–10499.

42. Stricker RL, Behrens SE, Mundt E: Nuclear factor NF45 interacts with viral proteins of infectious bursal disease virus and inhibits viral replication. J Virol 2010, 84(20):10592–10605.

43. Hammarlund E, Dasgupta A, Pinilla C, Norori P, Fruh K, Slifka MK: Monkeypox virus evades antiviral CD4+ and CD8+ T cell responses by suppressing cognate T cell activation. Proc Natl Acad Sci U S A 2008, 105(38):14567–14572.

44. Benfield CTO, Ren H, Lucas SJ, Bahsoun B, Smith GL: Vaccinia virus protein K7 is a virulence factor that alters the acute immune response to infection. J Gen Virol 2013, 94(Pt 7):1647–1657.

45. Torres AA, Macilwee SL, Rashid A, Cox SE, Albarnaz JD, Bonjardim CA, Smith GL: The actin nucleator Spir-1 is a virus restriction factor that promotes innate immune signalling. PLoS Pathog 2022, 18(2):e1010277.

46. Schroder M, Baran M, Bowie AG: Viral targeting of DEAD box protein 3 reveals its role in TBK1/IKKepsilon-mediated IRF activation. EMBO J 2008, 27(15):2147–2157.

47. Pichlmair A, Kandasamy K, Alvisi G, Mulhern O, Sacco R, Habjan M, Binder M, Stefanovic A, Eberle CA, Goncalves A et al: Viral immune modulators perturb the human molecular network by common and unique strategies. Nature 2012, 487(7408):486–490.

48. Zhou W, He Y, Rehman AU, Kong Y, Hong S, Ding G, Yalamanchili HK, Wan Y-W, Paul B, Wang C et al: Loss of function of NCOR1 and NCOR2 impairs memory through a novel GABAergic hypothalamus–CA3 projection. Nature Neuroscience 2019, 22(2):205–217.

49. Gileva IP, Nepomnyashchikh TS, Antonets DV, Lebedev LR, Kochneva GV, Grazhdantseva AV, Shchelkunov SN: Properties of the recombinant TNF-binding proteins from variola, monkeypox, and cowpox viruses are different. Biochim Biophys Acta 2006, 1764(11):1710–1718.

50. Alejo A, Ruiz-Arguello MB, Ho Y, Smith VP, Saraiva M, Alcami A: A chemokinebinding domain in the tumor necrosis factor receptor from variola (smallpox) virus. Proc Natl Acad Sci U S A 2006, 103(15):5995–6000.

51. Antonets DV, Nepomnyashchikh TS, Shchelkunov SN: SECRET domain of variola virus CrmB protein can be a member of poxviral type II chemokine-binding proteins family. BMC Res Notes 2010, 3:271.

