## Supplementary Table 2 for "Monkey Pox Virus (MPXV): Phylogenomics, Host-Pathogen Interactome, and Mutational Cascade"

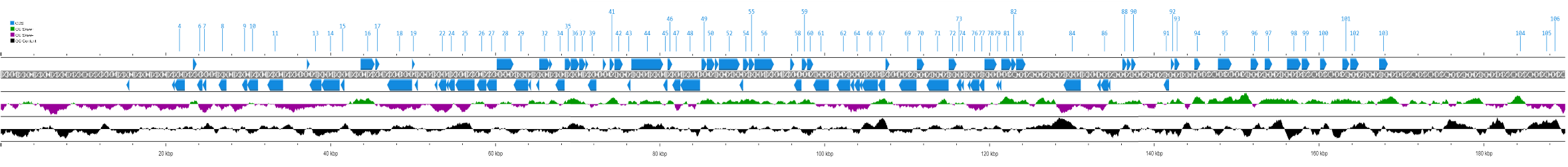

| **CDS  (Core Genome)** | **POSITION** | | **LENGTH (bp)** | **ACCESSION NUMBER** | **PROTEIN** | **PROTEIN ID** |
| --- | --- | --- | --- | --- | --- | --- |
| **1** | 15220 | 15573 | 354 | MN702448 | Anti-apoptotic Bcl-2-like protein | QNI39508 |
| **2** | 20694 | 20837 | 144 | MN702448 | Protein K3 | P20639 |
| **3** | 20841 | 20972 | 132 | MN702448 | IFN resistance PKR | QNI39514 |
| **4** | 21028 | 22302 | 1275 | MN702448 | Phospholipase-D-like protein | QNI39515 |
| **5** | 23296 | 23745 | 450 | MN702448 | Host immune response repressor | QNI39517 |
| **6** | 23789 | 24439 | 651 | MN702448 | Protein F1 | UniProtKB: P24356 |
| **7** | 24460 | 24894 | 435 | MN702448 | Deoxyuridine 5'-triphosphate nucleotidohydrolase | UniProtKB: P17374 |
| **8** | 26401 | 27360 | 960 | MN702448 | Ribonucleotide reductase small subunit | QNI39520 |
| **9** | 29243 | 29881 | 639 | MN702448 | S-S bond formation pathway protein | QNI39525 |
| **10** | 29868 | 31187 | 1320 | MN702448 | Ser-Thr kinase | QNI39526 |
| **11** | 32318 | 34225 | 1908 | MN702448 | EEV maturation protein | QNI39528 |
| **12** | 37136 | 37441 | 306 | MN702448 | DNA-binding virion core protein | QNI39534 |
| **13** | 37438 | 38877 | 1440 | MN702448 | Poly polymerase large subunit | QNI39535 |
| **14** | 38874 | 41087 | 2214 | MN702448 | IEV morphogenesis | QNI39536 |
| **15** | 41211 | 41672 | 462 | MN702448 | Double-stranded RNA-binding protein | QNI39537 |
| **16** | 43636 | 45339 | 1704 | MN702448 | Virion protein | QNI39539 |
| **17** | 45421 | 45921 | 501 | MN702448 | Putative myristoylated protein | QNI39540 |
| **18** | 46850 | 49870 | 3021 | MN702448 | DNA polymerase | QNI39542 |
| **19** | 49902 | 50189 | 288 | MN702448 | Sulfhydryl oxidase | QNI39543 |
| **20** | 50184 | 50573 | 390 | MN702448 | Putative virion core protein | QNI39544 |
| **21** | 52604 | 52930 | 327 | MN702448 | Glutaredoxin-1 | QNI39546 |
| **22** | 53077 | 54015 | 939 | MN702448 | Putative DNA-binding virion core protein | QNI39547 |
| **23** | 54022 | 54243 | 222 | MN702448 | IMV membrane protein | QNI39548 |
| **24** | 54244 | 55053 | 810 | MN702448 | DNA-binding phosphoprotein | QNI39549 |
| **25** | 55135 | 57450 | 2316 | MN702448 | Ribonucleotide reductase large subunit | QNI39550 |
| **26** | 57737 | 58885 | 1149 | MN702448 | Telomere binding protein | QNI39552 |
| **27** | 58878 | 60149 | 1272 | MN702448 | Virion core cysteine protease | QNI39553 |
| **28** | 60155 | 62185 | 2031 | MN702448 | RNA helicase NPH-II | QNI39554 |
| **29** | 62189 | 63964 | 1776 | MN702448 | Putative metalloprotease | QNI39555 |
| **30** | 63961 | 64200 | 240 | MN702448 | Entry/fusion complex component | QNI39556 |
| **31** | 64922 | 65296 | 375 | MN702448 | Glutaredoxin-like protein | QNI39558 |
| **32** | 65299 | 66603 | 1305 | MN702448 | FEN1-like nuclease | QNI39559 |
| **33** | 66612 | 66803 | 192 | MN702448 | RNA polymerase | QNI39560 |
| **34** | 67265 | 68380 | 1116 | MN702448 | Assembly Protein G7 | UniProtKB: P68715 |
| **35** | 68411 | 69193 | 783 | MN702448 | Late transcription factor VLTF-1 | QNI39563 |
| **36** | 69213 | 70235 | 1023 | MN702448 | Poxvirus myristoyl protein | QNI39564 |
| **37** | 70236 | 70988 | 753 | MN702448 | IMV membrane protein | QNI39565 |
| **38** | 71020 | 71298 | 279 | MN702448 | Crescent membrane and immature virion formation | QNI39566 |
| **39** | 71274 | 72308 | 1035 | MN702448 | Internal virion protein | QNI39567 |
| **40** | 73098 | 73484 | 387 | MN702448 | DNA-binding virion core protein | QNI39569 |
| **41** | 73919 | 74452 | 534 | MN702448 | Thymidine kinase | QNI39571 |
| **42** | 74518 | 75519 | 1002 | MN702448 | Poly polymerase small subunit | QNI39572 |
| **43** | 76051 | 76452 | 402 | MN702448 | Putative late 16 kDa membrane protein | QNI39574 |
| **44** | 76559 | 80419 | 3861 | MN702448 | RNA polymerase | QNI39575 |
| **45** | 80416 | 80931 | 516 | MN702448 | Tyr/Ser phosphatase, IFN-gamma inhibitor | QNI39576 |
| **46** | 80945 | 81514 | 510 | MN702448 | Putative viral membrane protein | QNI39577 |
| **47** | 81518 | 82492 | 975 | MN702448 | IMV heparin binding surface protein | QNI39578 |
| **48** | 82493 | 84880 | 2388 | MN702448 | RNA pol assoc protein | QNI39579 |
| **49** | 85065 | 85706 | 642 | MN702448 | Late transcription factor VLTF-4 | QNI39580 |
| **50** | 85707 | 86651 | 945 | MN702448 | DNA topoisomerase type I | QNI39581 |
| **51** | 86689 | 87129 | 441 | MN702448 | Crescent membrane and immature virion formation | QNI39582 |
| **52** | 87173 | 89710 | 2538 | MN702448 | mRNA capping enzyme large subunit | QNI39583 |
| **53** | 89669 | 89956 | 288 | MN702448 | Virion core | QNI39584 |
| **54** | 90102 | 90803 | 702 | MN702448 | Virion core | QNI39585 |
| **55** | 90803 | 91459 | 657 | MN702448 | Uracil-DNA glycosylase DNA polymerase | QNI39586 |
| **56** | 91491 | 93848 | 2358 | MN702448 | NTPase DNA primase | QNI39587 |
| **57** | 95828 | 96313 | 486 | MN702448 | DNA directed RNA Polymerase | QNI39589 |
| **58** | 96276 | 97190 | 915 | MN702448 | Carbonic anhydrase, GAG-binding IMV membrane protein | QNI39590 |
| **59** | 97232 | 97873 | 642 | MN702448 | mRNA decapping enzyme | QNI39591 |
| **60** | 97870 | 98616 | 747 | MN702448 | mRNA decapping protein | QNI39592 |
| **61** | 98617 | 100512 | 1896 | MN702448 | ATPase NPH1 | QNI39593 |
| **62** | 101441 | 103096 | 1656 | MN702448 | Trimeric virion coat protein | QNI39595 |
| **63** | 103121 | 103573 | 453 | MN702448 | Late transcription factor VLTF-2 | QNI39596 |
| **64** | 103594 | 104268 | 675 | MN702448 | Late transcription factor VLTF-3 | QNI39597 |
| **65** | 104265 | 104498 | 234 | MN702448 | S-S bond formation pathway protein | QNI39598 |
| **66** | 104513 | 106447 | 1935 | MN702448 | P4b precursor | QNI39599 |
| **67** | 106500 | 107345 | 846 | MN702448 | 39 kDa virion core protein | QNI39600 |
| **68** | 107383 | 107868 | 486 | MN702448 | RNA polymerase | QNI39601 |
| **69** | 109007 | 111139 | 2133 | MN702448 | Early transcription factor large | QNI39603 |
| **70** | 111193 | 112071 | 879 | MN702448 | Intermediate transcription factor 3 small subunit | UniProtKB: P20986 |
| **71** | 112355 | 115030 | 2676 | MN702448 | P4a precursor | QNI39606 |
| **72** | 115045 | 116001 | 957 | MN702448 | Viral membrane formation protein | QNI39607 |
| **73** | 116003 | 116575 | 573 | MN702448 | Virion core and cleavage processing protein | QNI39608 |
| **74** | 116599 | 116811 | 213 | MN702448 | IMV membrane protein | QNI39609 |
| **75** | 117357 | 117641 | 285 | MN702448 | Core Protein | QNI39612 |
| **76** | 117625 | 118758 | 1134 | MN702448 | Myristylated protein | QNI39613 |
| **77** | 118761 | 119375 | 615 | MN702448 | IMV membrane protein | QNI39614 |
| **78** | 119390 | 120868 | 1479 | MN702448 | DNA helicase | QNI39615 |
| **79** | 120849 | 121082 | 234 | MN702448 | Zinc finger-like protein | QNI39616 |
| **80** | 121083 | 121355 | 273 | MN702448 | IMV membrane protein | QNI39617 |
| **81** | 121429 | 122709 | 1281 | MN702448 | DNA polymerase processivity factor | QNI39618 |
| **82** | 122639 | 123202 | 519 | MN702448 | Holliday junction resolvase | QNI39619 |
| **83** | 123222 | 124370 | 1149 | MN702448 | VITF-3 45 kDa subunit | QNI39620 |
| **84** | 128975 | 131065 | 2091 | MN702448 | A-type inclusion protein | QNI39623 |
| **85** | 133045 | 133485 | 441 | MN702448 | IMV MP virus entry protein | QNI39625 |
| **86** | 133486 | 134403 | 918 | MN702448 | DNA polymerase | QNI39626 |
| **87** | 134366 | 134599 | 234 | MN702448 | IMV Protein | QNI39627 |
| **88** | 136102 | 136647 | 546 | MN702448 | EEV membrane phosphoglycoprotein C-type | QNI39631 |
| **89** | 136652 | 137158 | 507 | MN702448 | IEV and EEV membrane glycoprotein | QNI39632 |
| **90** | 137202 | 137732 | 531 | MN702448 | MHC class II antigen presentation inhibitor | QNI39633 |
| **91** | 141134 | 141775 | 642 | MN702448 | Chemokine binding protein | QNI39638 |
| **92** | 142017 | 142418 | 402 | MN702448 | Profilin-like protein ATI-localized | QNI39639 |
| **93** | 142462 | 143061 | 600 | MN702448 | Type I membrane glycoprotein | QNI39640 |
| **94** | 144856 | 145578 | 723 | MN702448 | Toll/IL1-receptor [TIR]-like protein | QNI39644 |
| **95** | 147725 | 149492 | 1768 | MN702448 | ATP-dependent DNA ligase | QNI39646 |
| **96** | 151693 | 152634 | 942 | MN702448 | Hemagglutinin | QNI39649 |
| **97** | 153401 | 154312 | 912 | MN702448 | Serine-Threonine kinase | QNI39650 |
| **98** | 156104 | 157789 | 1695 | MN702448 | Ankyrin repeat domain-containing protein M-T5 | UniProtKB: Q83730 |
| **99** | 157893 | 158846 | 954 | MN702448 | EEV membrane glycoprotein | QNI39653 |
| **100** | 160116 | 160919 | 804 | MN702448 | Soluble interferon-gamma receptor-like protein | QNI39656 |
| **101** | 162820 | 163668 | 849 | MN702448 | Ser-Thr kinase | QNI39659 |
| **102** | 163776 | 164810 | 1035 | MN702448 | Ser-Thr kinase | QNI39660 |
| **103** | 167278 | 168340 | 1063 | MN702448 | IFN-alpha/beta receptor glycoprotein | QNI39663 |
| **104** | 183752 | 185065 | 1314 | MN702448 | Ankyrin repeat domain | UniProtKB: [Q86XL3](https://www.uniprot.org/uniprotkb/Q86XL3/entry) |
| **105** | 187075 | 188121 | 1047 | MN702448 | Ankyrin repeat domain | UniProtKB: [Q86XL3](https://www.uniprot.org/uniprotkb/Q86XL3/entry) |
| **106** | 188249 | 189007 | 759 | MN702448 | Chemokine binding protein | UniProtKB: [O00590](https://www.uniprot.org/uniprotkb/O00590/entry) |
